# A micro-PRNT for the detection of Ross River virus antibodies in mosquito blood meals: a useful tool for inferring transmission pathways

**DOI:** 10.1101/2020.02.05.935437

**Authors:** Narayan Gyawali, Amanda Murphy, Leon E. Hugo, Gregor J Devine

## Abstract

**Introduction:** Many arboviruses of public health significance are maintained in zoonotic cycles with complex transmission pathways. The presence of serum antibody against arboviruses in vertebrates provides evidence of their historical exposure but reveals nothing about the vector-reservoir relationship. Moreover, collecting blood or tissue samples from vertebrate hosts is ethically and logistically challenging. We developed a novel approach for screening the immune status of vertebrates against Ross River virus that allows us to implicate the vectors that form the transmission pathway for this commonly notified Australian arboviral disease.

**Methods:** A micro-plaque reduction neutralisation test (micro-PRNT) was developed and validated on koala (*Phascolarctos cinereus)* sera against a standard PRNT. The ability of the micro-PRNT to detect RRV antibodies in mosquito blood meals was then tested using some convenient mosquito models. Laboratory-reared *Aedes aegypti* were fed, via a membrane, on sheep blood supplemented with RRV seropositive and seronegative human sera. *Aedes notoscriptus* were fed on RRV seropositive and seronegative human volunteers. Blood-fed mosquitoes were harvested at various time points after feeding and their blood meals analysed for the presence of RRV neutralizing antibodies using the micro-PRNT.

**Results:** There was significant agreement of the plaque neutralization resulting from the micro-PRNT and standard PRNT techniques (R^2^=0.65; P<0.0001) when applied to RRV antibody detection in koala sera. Sensitivity and specificity of the micro-PRNT assay were 88.2% and 96%, respectively, in comparison with the standard PRNT. Blood meals from mosquitoes fed on sheep blood supplemented with RRV antibodies neutralised RRV by ≥50% until 60 hr post feeding. Similarly, mosquito blood meals from RRV seropositive human volunteers neutralised the virus by ≥50% until 48 hr post-blood feeding.

**Conclusions:** The small volumes of blood present in mosquito abdomens can be used to identify RRV antibodies and therefore host exposure to arbovirus infection. In tandem with the accurate identification of the mosquito, and diagnostics for the host origin of the blood meal, this technique has tremendous potential for exploring RRV transmission pathways. It can clearly be adapted for similar studies on other mosquito borne zoonoses.

## Introduction

Arthropod borne viruses (arboviruses) present a significant risk to public health globally. In recent decades, rapid urbanization and population growth have assisted the development of several viruses from having a localised, rural, transmission cycles to becoming worldwide and urban problems [1]. Epidemiological cycles of many arboviruses, such as Ross River (RRV) and West Nile, incorporate complex transmission networks involving multiple vertebrate hosts and many vectors. Humans are not necessarily key components of the transmission network, but increasing human travel, trade and deforestation bring humans into contact with sylvatic/enzootic cycles. This can stimulate arbovirus emergence, re-emergence and spillover into human populations [2-4].

A comprehensive knowledge of the transmission pathways of arboviruses is needed to effectively manage and respond to their emergence. Surveillance systems are needed to identify which mosquito species are responsible for transmission and which animals are acting as amplifying or reservoir hosts. However, the identification of amplifying hosts and transmission pathways remains extremely challenging.

More than 75 arboviruses have been identified in Australia and a small number are associated with human infection [5]. Of these, RRV [6], Barmah Forest virus [7], West Nile virus strain Kunjin [8], and the potentially fatal Murray Valley encephalitis virus [9] are of the greatest public health concern. RRV is the most commonly notified arboviral disease but multiple vectors and many potential vertebrate hosts make this a complex zoonosis. There is little empirical evidence regarding its transmission cycles or encourage their spillover to the human population [10, 11].

One means of identifying likely vertebrate disease reservoirs is to demonstrate their historical exposure to disease by searching for virus-specific antibodies in animal sera or tissues. Development of antibody is the major immune response to infection with parasites and pathogens including the arboviruses [12, 13]. While such serological evidence of infection does not prove that an animal is an amplifying host or key reservoir, it does allow us to generate hypotheses about probable pathways and is especially useful when combined with information on mosquito species and their host preference. Serological surveys of blood meals are likely to be more fruitful than the direct identification of viruses because vertebrates are only viraemic for a few days, only a small proportion of mosquitoes are virus positive and there is a diminishingly small probability that a mosquito will be carrying a virus positive mosquito blood meal. A tremendous sampling effort is therefore required to incriminate reservoir and vector pathways by virus isolation alone.

In our alternative approach, we exploit the fact that vertebrate antibodies persist within mosquito blood meals for some time after the mosquito has fed on a seropositive host. We demonstrate that a micro-PRNT technique can identify vertebrate RRV antibodies in small volumes of sera and mosquito blood meals. This has utility as part of an integrated xenodiagnostic approach that exploits the capture of single blood-fed mosquitoes to infer mosquito species, host preference and host exposure to disease. This will help prioritise potential transmission pathways for further study.

The potential for screening mosquito blood meals for antibodies to dengue, Japanese encephalitis [14], and West Nile Virus [15] has been investigated previously but existing studies required the use of host-specific conjugated antibodies. This is of little utility for the investigation of complex zoonoses like RRV where the hosts are myriad or unknown.

The “gold standard” of serological tests is the Plaque Reduction Neutralization Test (PRNT)[16]. It does not need prior knowledge of host origin but typically requires large quantities of sera or tissue; substantially larger than a typical mosquito blood meal (ca 3 µl [17]). We developed a micro-PRNT [18, 19] to suit small sample volumes.

## Methods

### Cells and virus

Vero cells (WHO vaccine strain) and the RRV strain T-48 [20] were obtained from the WHO Collaborating Centre for Arbovirus Reference and Research at the Queensland University of Technology. At the QIMR Berghofer Medical Research Institute (QIMRB), virus was propagated in Vero cells maintained in 5% CO_2_ at 37°C in RPMI-1640 growth media (Sigma-Aldrich, Missouri, USA), supplemented with L-glutamine (0.3 g/L), sodium bicarbonate (2 g/L), 10% (v/v) heat-inactivated foetal calf serum (Invitrogen, USA) and 1% (v/v) PSG [(Penicillin (10,000 units)/Streptomycin (10 mg/mL)/ L-glutamine (200 mM)); Sigma-Aldrich, USA]. Virus stocks were frozen at −80°C until further use.

### Koala sera

Forty-two koala sera, obtained from Endeavour Vets, Queensland, Australia (http://www.endeavourvet.com.au) were used to validate the micro-PRNT. These samples were collected between 2015 to 2017 and stored at −80 °C.

### Mosquitoes

We used two model insects to validate the micro-PRNT. An *Ae. aegypti* colony originated from Cairns, Australia, in 2015 and was reared as previously described [21]. Adult mosquitoes were provided with 10% sugar solution *ad libitum* and an opportunity to feed on defibrinated sheep blood once per week. An *Ae. notoscriptus* colony was established from eggs collected in Brisbane, Australia during 2015 and maintained as above.

### Development of the microPRNT

All koala serum samples were tested for neutralising RRV antibodies using a conventional PRNT approach. Equal volumes of sera (200 µl), diluted 1:20 in serum-free RPMI-1640, were mixed with an equal volume of 50 plaque forming unit RRV (1:800 stock RRV in RPMI-1640) per well of a 12-well tissue culture plate (Nunclon, Thermo Scientific, Australia). The virus-sera mixtures were incubated at 37°C for 45 min and added to infect Vero cell monolayers and incubated for a further two hours to enable non-neutralised virus to adsorb to cells. Following incubation, the virus-sera mixture was removed and 2 mL of 0.75% w/v carboxymethyl cellulose (CMC, Sigma-Aldrich) /RPMI 1640 was added. Plates were incubated at 37°C in 5% v/v CO_2_ for an additional 40 hr. The CMC/RPMI medium was then removed, and the cell monolayers were fixed and stained with 0.05% w/v crystal violet (Sigma-Aldrich) in formaldehyde (1% v/v) and methanol (1% v/v). The cell monolayers were then rinsed in tap water, and the plates inverted on a paper towel until dry. Plaques (clear zones in a purple cell monolayer) were counted. Reductions of total virus plaque numbers per well of ≥50% were considered to denote seropositive status [13].

All koala samples were also tested by a micro-PRNT technique using just three µl of koala sera; the approximate volume of a mosquito blood meal [17], also diluted 1:20 in serum-free RPMI-1640. Equal volumes of sera (50 µl) were mixed with an equal volume of 30 pfu RRV (1:160 stock RRV in RPMI-1640) and added to wells of 96–well tissue culture plate (Nunclon, Thermo Scientific, Australia) containing a Vero cell monolayer. Thirty plaques per well in 96-well plates are easier to discriminate given the small field of view [22]. A volume of 200µl CMC/RPMI was added to each well following infection of the cell monolayer.

The agreement between the percent plaque neutralisation from both the conventional and micro-PRNT was determined by paired sample t-test (n=42, each serum sample tested once with each PRNT protocol).

### Preparation of blood fed mosquitoes

Defibrinated sheep blood (Serum Australis, Manilla, NSW, Australia) was confirmed seronegative for RRV by PRNT and washed three times with saline. Two sets of washed-sheep blood cells were supplemented with RRV seropositive and RRV seronegative human sera from human volunteers (QIMR Berghofer ethics ref: P2273) at a proportion of 55 to 45 to keep the composition of plasma and blood cells realistic. Female *Ae. aegypti* mosquitoes (aged 3-5 days) were starved of sugar solution for 5 hrs to increase their avidity and then offered seropositive or seronegative blood for 30-45 min via a membrane feeding apparatus [23]. Fully engorged and unfed mosquitoes from these treatments were maintained separately at 27±1°C and 80% RH humidity and provided with 10% sucrose solution *ad libitum*. At 6, 12, 24, 36, 48, 60 and 72 hr post exposure to the infected and uninfected blood meals, fed and unfed mosquitoes (the latter used as controls) were harvested and stored at −80 °C.

In a separate experiment, to fully validate the micro-PRNT on blood meals from mosquitoes that had fed directly on vertebrate hosts, *Ae. notoscriptus* were fed on the exposed arms of RRV seropositive and seronegative human volunteers (confirmed by PRNT) for a period of 15 min (QIMR Berghofer ethics ref: P2273). Blood-fed mosquitoes were maintained as described above and harvested at 6, 12, 24, 36, 48, 60 and 72 hr post feeding. Remnant blood meals could be observed in mosquito abdomens until 60 hours (Fig 1).

**Fig 1.**
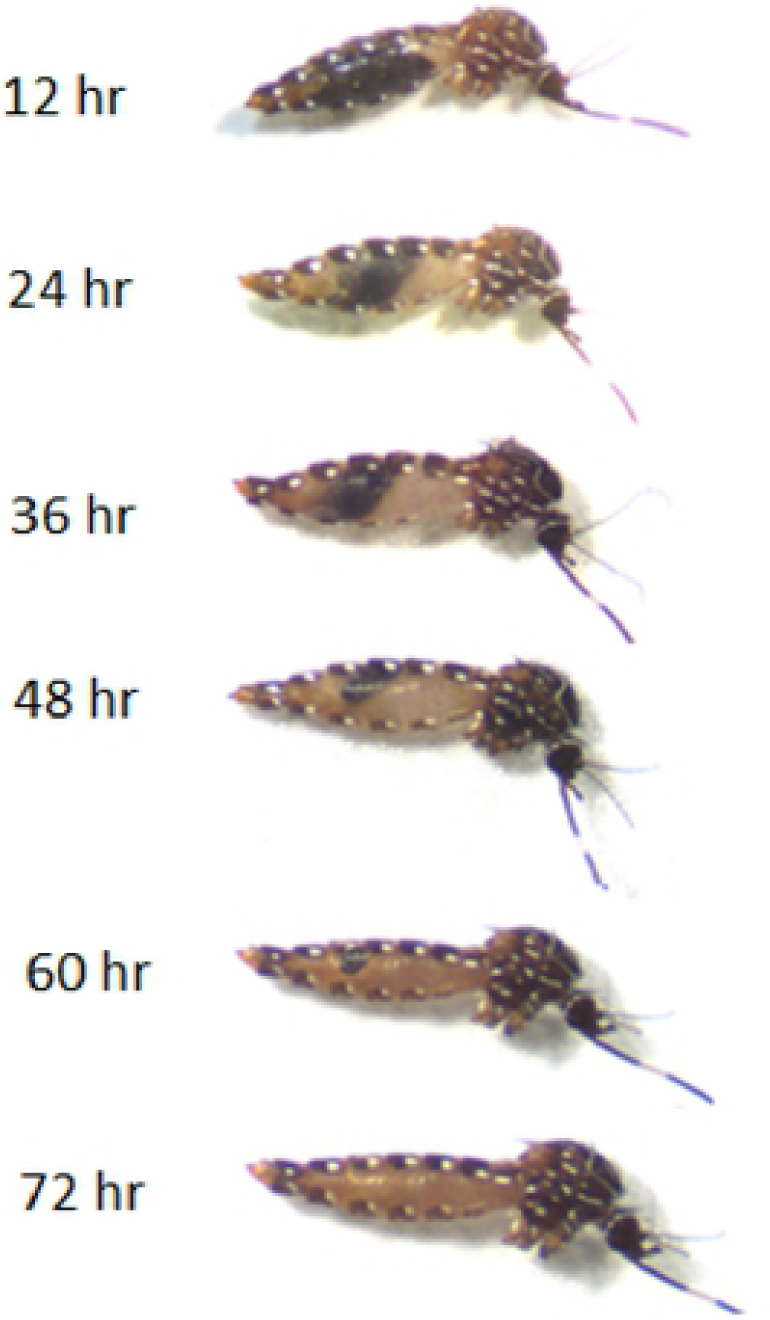
*Aedes notoscriptus* at different times during blood meal digestion.

### Validation of the micro-PRNT using blood-fed mosquitoes

The blood meal volume obtained from a single field-collected, blood-fed mosquito is sufficient to test in the micro-PRNT assay, but in order to ascertain the assay’s robustness against a range of host antibody titres and post-feeding times, and facilitate the dilutions that these experiments required, we also used larger volumes of mosquito-derived blood in our validations. These larger volumes were obtained by pooling three blood meals from engorged mosquitoes that had fed on the same antibody-positive source. One pool was used for each post-feeding time point tested. Abdominal contents were expelled into serum-free RPMI 1640 supplemented with 1% PSG and 0.4% amphotericin B (Sigma-Aldrich, USA). The volume of RPMI 1640 was adjusted to obtain a 20-fold dilution of the blood. Each sample was centrifuged at 10,000 × g for 10 min and the supernatant was transferred to a sterile Eppendorf tube and further dilutions of 1:40, 1:80 and 1:160 were derived for each sample. Positive and negative controls were included in each assay batch to validate the technique and to assess the impacts of mosquito homogenates alone on the formation of plaque forming units: 1) RRV alone 2) RRV plus unfed mosquito abdomens, and 3) RRV plus mosquitoes fed with sheep blood supplemented with RRV seronegative human sera. In every case, the pfu RRV was kept constant.

The final proof that the RRV micro-PRNT can be used on single, blood fed mosquitoes was conducted using *Aedes notoscriptus* that had been fed on human seropositive or seronegative volunteers. Blood-fed mosquitoes were harvested at various times post-feeding and processed as above (abdominal contents expelled into serum-free RPMI 1640 and adjusted to obtain a 20-fold dilution). Unfed mosquitoes were included as a control.

### Plaque counting

Plates were scanned at 600 dpi resolution (HP Scanjet, Palo Alto, USA) and images were magnified before counting plaques.

### Host identification by molecular method

Sixteen blood meal samples (equal numbers, 8 of each) harvested after feeding on RRV seropositive sheep blood or human blood were used to prove the principal of parallel host identification using Cytochrome b DNA amplified by established PCR techniques [22]. Ten µl of the blood meal homogenate remaining after the micro PRNT was used for blood-meal host identification (DNA extraction, amplification and sequencing) using published methods [24].

## Results

### Comparison of micro-PRNT and PRNT

RRV plaques on cell layers stained 40 hr post-incubation were clearly distinguishable and easily counted when plates were scanned and images enlarged. There was a significant correlation (R^2^=0.65; P<0.0001) between percent neutralization of RRV pfu noted in koala samples characterized by micro-PRNT or PRNT techniques (Fig 2). In comparison to the standard PRNT, the sensitivity and specificity of the micro-PRNT was 88.2% and 96% respectively. Those differences in percent neutralisation between methods were not significant (*p* > 0.05; paired samples t-test).

**Fig 2.**
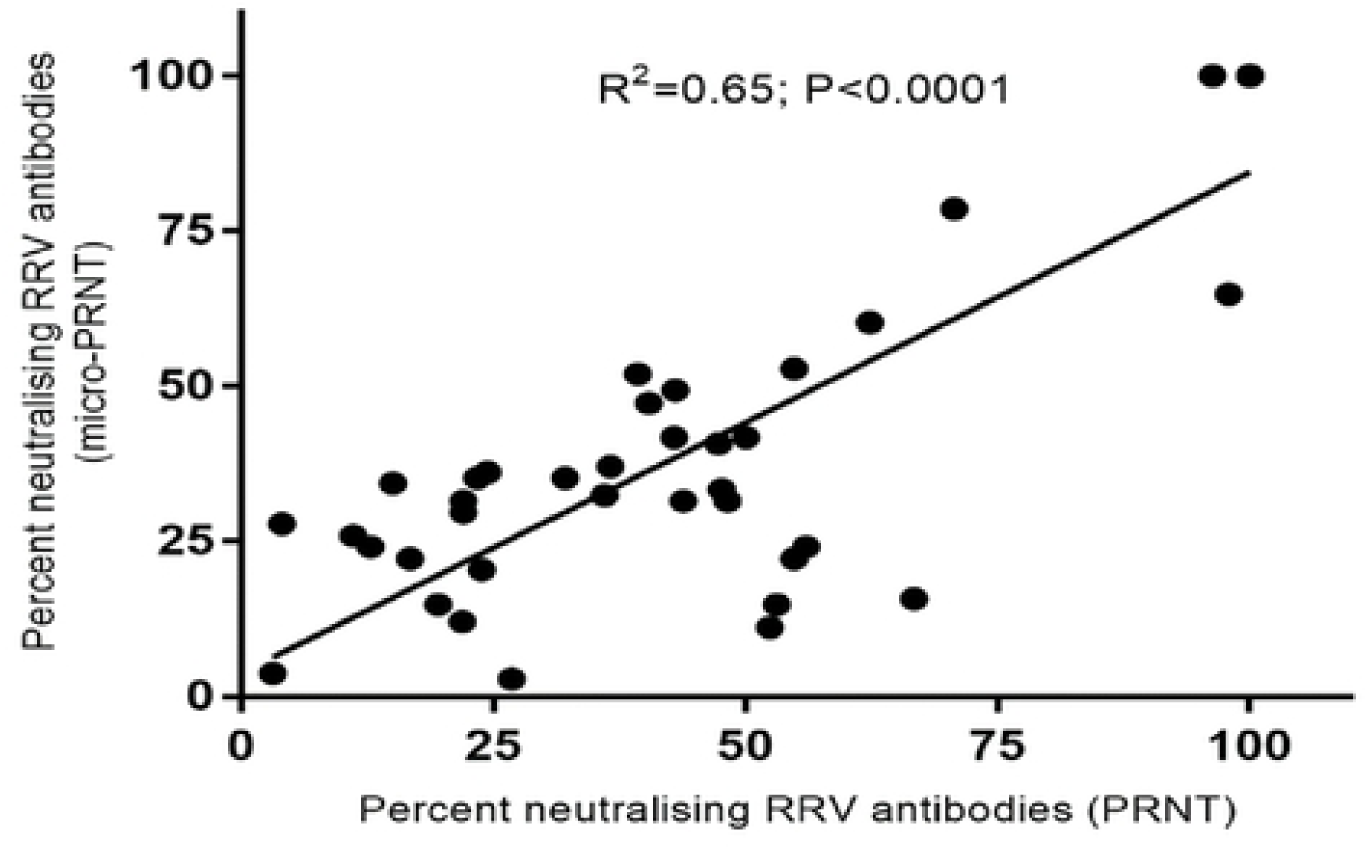
Agreement between RRV percent neutralising antibodies determined by standard PRNT or micro-PRNT using koala serum samples.

### Impacts of mosquito homogenates on the number of plaque forming units

Vero cells were infected by RRV alone, RRV mixed with homogenates of unfed mosquito abdomens or abdomens containing RRV seronegative sheep blood. There was a small (10%) but significant decrease in RRV pfu when Vero cells in 96 well were infected by RRV mixed with the latter two samples (Fig 3).

**Fig 3.**
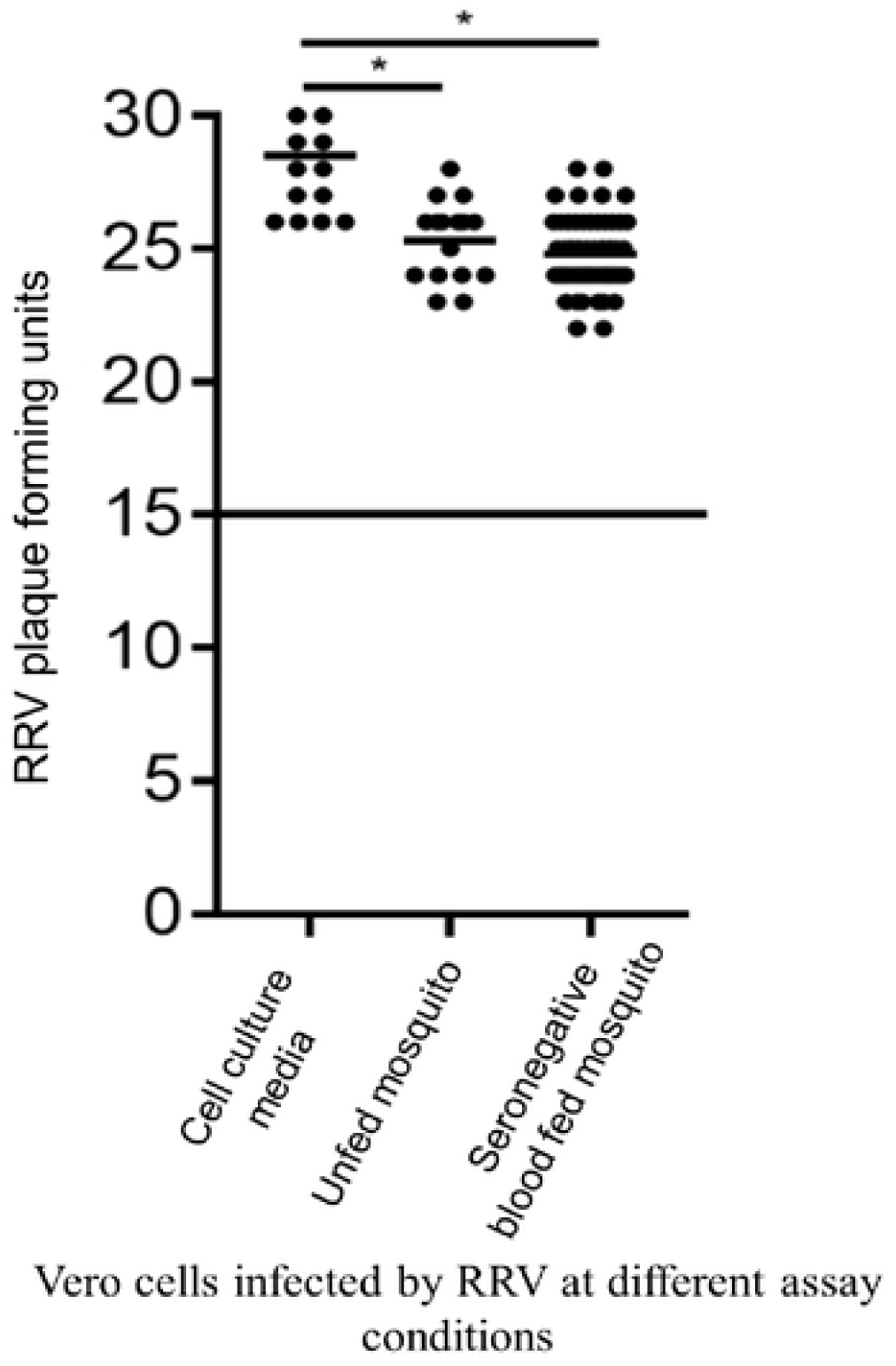
The effect of mosquito homogenates on plaque formation. *p<0.05, calculated by one-way ANOVA test.

### Validation of the assay using mosquito blood meals

Mosquito blood meals obtained from mosquitoes fed on sheep blood mixed with anti-RRV human antibodies neutralised RRV by ≥50% until 60 hr post blood feeding (Figs 4a and 4b). In contrast, there was no neutralization of blood meals obtained from mosquitoes fed with sheep blood supplemented with RRV seronegative human serum (Fig 4c). These assays demonstrated that the micro-PRNT is robust across a range of dilutions that is likely to represent antibody levels in different hosts (Fig 4a).

**Fig 4.**
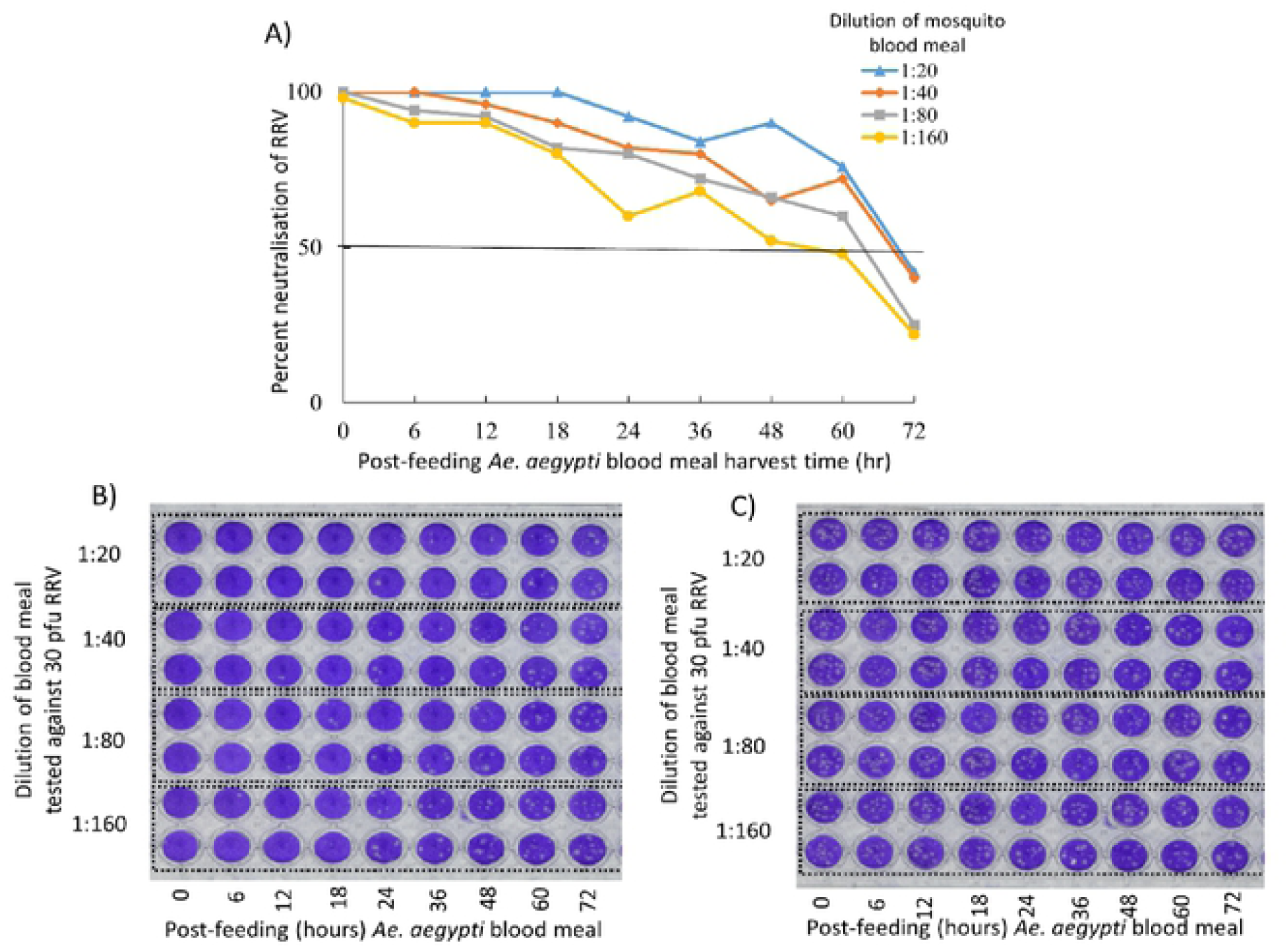
Percent neutralisation of RRV by mosquito blood meals harvested at different time points post blood-feeding (A). Appearance of stained cell monolayers following incubation with RRV and mosquito abdomens containing RRV seropositive sera (B) and RRV seronegative sera (C). Samples were plated in duplicate for each dilution (rows) and for each period post-feeding (columns). Plaques are evident as unstained regions.

Guided by the results detailed in Fig 4, we used the 1:20 dilution for all micro-PRNTs on single blood meals from live hosts. Single *Ae. notoscriptus* blood meals obtained through feeding mosquitoes on a seropositive human volunteer were harvested at different time points. These blood meal preparations neutralised RRV pfu by ≥50% until 36-48 hr post feeding. (Fig 5A). The impact of sero-negative blood meals (Fig 5B) and of abdomens from un-fed mosquitoes are included for comparison (Fig 5C)

**Fig 5.**
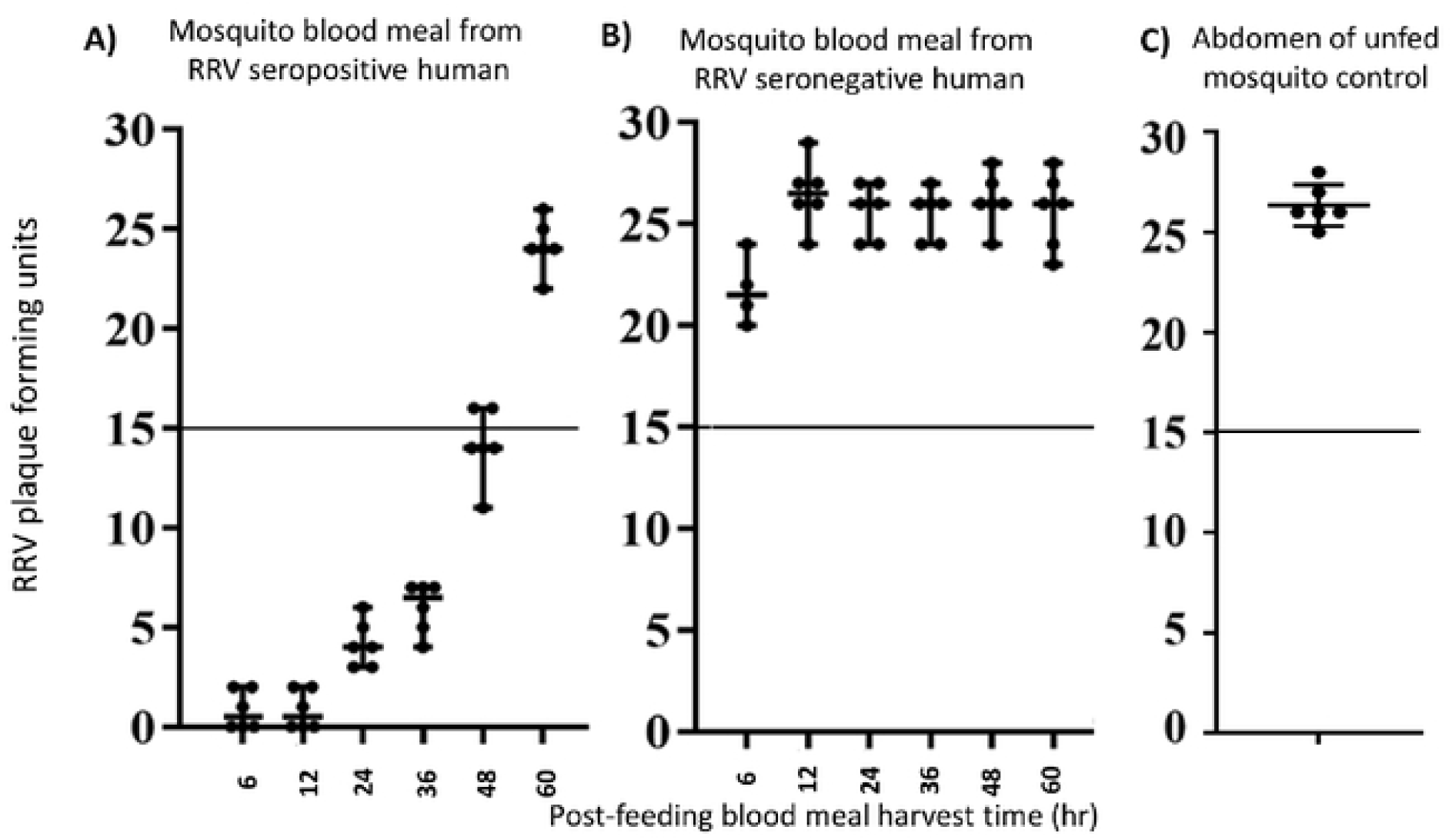
RRV micro-PRNT with *Aedes notoscriptus* blood meals harvested at various times post-feeding. blood meals from an RRV seropositive human volunteer (A), blood meals from an RRV seronegative human volunteer (B) and unfed mosquito abdomens as a control (C).

### PCR of blood meal samples

Sequencing of amplicon-DNA from all 16 samples correctly identified the origin of the blood meal source (sheep blood from those experiments that had used membrane feeds, and human where mosquitoes had fed on a live host

## Discussion

Mosquito blood meals are a potentially useful resource for assessing antibody seroprevalence in vertebrates and inferring the transmission pathways between vectors, disease reservoirs and humans. This study demonstrates that a micro-PRNT using 96-well plates has considerable utility for characterizing the antibody content of small blood volumes. In conjunction with the species identification of the blood-fed mosquito, and the use of existing tools to identify the host origin of the blood meal, this new diagnostic has the capacity to increase our understanding of the key pathways for the transmission of complex zoonotic arboviruses.

The PRNT is the “gold standard” tool for identifying neutralizing antibodies in vertebrate blood and tissue samples. The accuracy of the micro-PRNT for testing RRV antibodies in this study was similar to that reported for the micro-PRNT developed for testing yellow fever antibodies [18]. However, these results from both, Simoes et al. (2012) and the current (sero-detection of RRV in koala sera) studies were not from mosquito blood meal.

Alternatives to the PRNT such as a VecTest-inhibition assay and a biotin microsphere immunoassay have been used to identify pathogen specific antibodies in mosquito blood meals [15], but both require host specific antibody and the latter demands considerable laboratory resources. The micro-PRNT demonstrates sufficient sensitivity, specificity and utility for the determination of RRV antibodies in mosquito blood meals.

Our micro-PRNT could detect ≥50% RRV neutralization in mosquito blood meals up to 36-48 hr post blood-feeding. Although this is the first study to have identified RRV antibodies in mosquito blood meals, Hatfield (1988) identified Bovine Serum Albumin (BSA) specific antibodies using antibody-captured ELISA from the hemolymph of *Ae. aegypti* up to 48 hr after feeding [25]. Irby and Apperson (1989) used an immunoblot technique to demonstrate that serum proteins such as albumin and immunoglobulin G of rodents and human persisted in mosquito (*Ae. aegypti*) blood meal for 36 to 48 hr of post-feeding [26]. In other studies this detection period was far greater: using ELISA methods, anti-BSA antibodies were detected 9 days after blood feeding in *Anopheles stephensi* [27] and human specific IgM and IgG were present in the blood meals of *Ae. albopictus* for 7 days [28]. This is surprising given that one would expect blood meals and their proteins to have been fully digested by then but the persistence of antibodies in mosquito blood meals will differ with species, ambient temperature, body size, initial concentration of antibodies, blood meal volume and the length of the gonotrophic cycle. In our study, the period of detectability (48-60 hr) corresponded to the period that blood was externally visible in the abdomen (Fig 1).

The small volumes of homogenate left from a single mosquito from the micro-PRNT allows for a parallel PCR amplification for identification of the blood meal source (the host). The literature commonly reports that the origin of host blood meals can be identified from as a little as 0.02 µl of blood [29]. In our proof of principle, 10 ul aliquots of 1:20 diluted homogenate recovered from the micro-PRNT 60 hr post blood-feeding were successfully amplified and sequenced to identify our experimental donors: sheep and humans.

Given the challenges involved in obtaining, trapping and screening wild animals for serum sampling and sero-prevalence studies [13], the collection of blood fed mosquitoes is an useful means of sampling, with mosquitoes acting as an indirect sampling tool or “flying syringe” for sampling inaccessible or ethically challenging hosts. In terms of pathway incrimination, a single mosquito will yield information on vectors, host preference and the disease exposure of that host [11]. That information, especially when combined with risk modelling [30] can be used to identify those transmission pathways of greatest importance within the host community. Various studies have observed the potential for insect blood meals to determine pathogen specific antibodies [25, 31-33], but none have developed high throughput methods suitable for application to transmission ecologies involving unknown reservoirs.

Human health can be affected by infectious diseases of wildlife living close to human habitation. The risks are increasingly common in Australia and elsewhere because increasing encroachment of the human population on diverse mosquito habitats and the potential adaptation of reservoir species to urbanized environments [34, 35]. Dengue, Hantavirus, Lyme disease, Zika, avian influenza, and rabies are examples of globally endemic zoonoses that have emerged from human encroachment into rural or sylvatic habitats [36, 37]. Similarly, RRV is a major public health risk in Australia, maintained in a diverse range of hosts and vectors and undergoing an expansion in range to the Pacific Islands [38].

Infectious diseases are also a concern for wildlife conservation, particularly those already threatened by habitat loss and exploitation. Surveillance of wild animals for infection or disease commonly involves trapping or killing animals for direct sampling of blood and tissues. This can be difficult, expensive, dangerous and sometime unethical. The technique demonstrated here is not only applicable to RRV reservoir identification but also to other arboviruses and infectious agents which have complex transmission cycles and a range of non-human vertebrate hosts.

There was ca 10% inhibition of virus pfu by mosquito tissue homogenates (Fig 3). However this inhibition was far less the threshold used to determine positive neutralization (>50% reduction). One possible explanation with this observation is that, when crushing mosquito abdomens, the sample might include some abdominal tissue that inhibits virus replication.

As for all serum or tissue collection techniques, reliance on blood-fed mosquitoes as a sampling tool will be subject to sampling bias. Mosquitoes may be differentially attracted to diseased hosts [39] or to species that are uncharacteristically abundant at any single point in time. Different mosquito trapping techniques have differential vector specific targets, so using one particular trap type might miss key vector species. In terms of serology, there may be considerable cross reaction between some virus antibodies [12]. In this case RRV is likely to cross-react with the closely related alphavirus, Barmah Forest virus, whose epidemiology in Queensland remains significant [2]. Finally, those reservoirs inferred by blood meal analysis may not be the key amplifying host, and the mosquitoes incriminated may not be the key vector. For example, dengue antibodies are commonly found in *Culex* spp. mosquitoes during epidemics, but those mosquitoes do not transmit the disease. Nonetheless their blood meals may still identify the host and its sero-prevalence rate [14]. For all of these reasons, the various components of the pathways implicated by the micro-PRNT technique and attendant host identifications must be interpreted and prioritised in the light of all the available knowledge on the ecology of the disease.

## Conclusion

The application of the micro-PRNT technique to vertebrate blood samples sourced from mosquito blood meals presents a novel xenodiagnostic for determining the sero-epidemiology of arboviruses and to infer transmission pathways for further investigation. It offers an alternative “flying syringe” approach for serum sampling and for monitoring sero-prevalence in animals. The use of this assay to characterise the blood meals of mosquitoes collected from the field in Brisbane is in preparation.

## Funding and acknowledgement

This work was supported by an Australian Post Graduate Award to Amanda Murphy, by funds from the Mosquito Control Laboratory, QIMR Berghofer and by the Mosquito and Arbovirus Research Committee (MARC) which is an independent Australian organization funded by local government, government agencies, industry and scientific institutions. We thank John Aaskov and Francesca Frentiu (Queensland University of Technology) for their advice and for the supply of vero cells and the RRV isolate.

## Authors’ Contributions

NG, AM, LEH and GD conceived the project. NG, AM and LEH carried out the laboratory experiments. NG drafted the paper. GD, LEH, AM reviewed and revised the manuscript. All authors contributed to preparation of the final version and agreed to its submission.

